# Mosquito bites and stages specific antibody responses against *Plasmodium falciparum* in southern Ghana

**DOI:** 10.1101/2022.11.03.514987

**Authors:** Sebastian Shine Kwapong, Kwame Kumi Asare, Kwadwo Asamoah Kusi, Faustina Pappoe, Nicaise Ndam, Rachida Tahar, Anne Poinsignon, Linda Eva Amoah

**Affiliations:** Department of Microbiology and Immunology, School of Medical Sciences, University of Cape Coast, Cape Coast, Ghana; Department of Biomedical Science, School of Allied Health Sciences, University of Cape Coast, Cape Coast, Ghana; Department of Immunology, Noguchi Memorial Institute for Medical Research, University of Ghana, Accra, Ghana; Department of Parasitology, Noguchi Memorial Institute for Medical Research, University of Ghana, Accra, Ghana; UMR261 MERIT, Accra, Ghana; Université de Paris Cité, MERIT, IRD, 75006 Paris, France; MIVEGEC, University of Montpellier, IRD, CNRS, 34000, Montpellier, France; Biomedical and Clinical Research Centre, College of Allied Health Sciences, University of Cape Coast, Cape Coast, Ghana

## Abstract

**Background:** The human host elicits specific immune responses after exposure to various life stages of the malaria parasite as well as components of mosquito saliva injected into the host during a mosquito bite. This study describes differences in IgG responses against antigens derived from the sporozoite (PfCSP), asexual stage parasite (PfEBA175) and the gametocyte (Pfs230) in addition to an *Anopheles gambiae* salivary gland antigen (gSG6-P1) in two communities in Ghana with similar blood stage malaria parasite prevalence.

**Methodology:** This study used archived plasma samples collected from an earlier cross-sectional study that enrolled volunteers aged from 6 months to 70 years from Simiw, peri-urban community (N=347) and Obom, rural community (N=291). An archived thick and thin blood smear for microscopy was used for the estimation of *Plasmodium* parasite density and species and DNA extraction from blood spots and *P. falciparum* confirmation was performed using PCR. This study used the stored plasma samples to determine IgG antibody levels to *Plasmodium falciparum* and *Anopheles* salivary antigens using indirect ELISA.

**Results:** Individuals from Simiw had significantly higher levels of IgG against mosquito gSG6-P1 (median (95%CI)) (2.590 (2.452-2.783) ng/mL) compared to those from Obom (2.119 (1.957-2.345) ng/mL), p<0.0001. Both IgG responses against Pfs230proC (p=0.0006), and PfCSP (p=0.002) were significantly lower in volunteers from Simiw compared to the participants from Obom. The seroprevalence of PfEBA-175.5R (p=0.8613), gSG6-P1 (p=0.0704), PfCSP (p=0.7798) IgG were all similar in Obom and Simiw. However, Pfs230 seroprevalence was significantly higher at Obom compared to Simiw (p=0.0006). Spearman correlation analysis showed no significant association between IgG responses against gSG6-P1, PfCSP, Pfs230proC and PfEBA-175.5R and parasite density at both Obom and Simiw (p>0.05).

**Conclusion:** In conclusion, malaria transmission dynamics is highly complex. The similar malaria transmission intensity identified in the two communities resulted from a different combination of vector, environmental and host factors. With one community likely having a higher prevalence of uninfected mosquitoes and the other a larger reservoir of gametocyte carriers.

## Introduction

Sustaining the gains in malaria control requires efforts to develop new and sensitive diagnostic tools, interventions to reduce contact with the mosquito vector as well as new drugs and vaccines. *Plasmodium* parasites undergo a complex life cycle within the mosquito vector and the human host [1, 2]. Exposure to various life cycle stages of the malaria parasite results in the host eliciting immune responses that include specific antibody responses against these different parasite life cycle stages. Antibody responses against the malaria parasite can be indicative of exposure and thus serve as markers of infection or serve a specific function such as used to regulate the infection or block transmission [3].

*Plasmodium falciparum* infections containing gametocytes can result in the development of transmission blocking immunity, which prevents the completion of the parasite’s life cycle in the mosquito as well as antibodies that are indicative of exposure to gametocytes. Mosquitoes pick-up gametocytes develop into sporozoites which subsequent infect the human host during a blood meal. Sporozoites are the first parasite stage to be encountered by the human host and initiate an infection. The surface of sporozoites has antigens including the circumsporozoite protein (PfCSP) that induces cell-mediated and antibody immune responses in the host [4, 5]. Immune responses generated by a host against this parasite stage prevents the establishment of an infection within the liver [6].

*Plasmodium falciparum* merozoites develop from sporozoites and invade erythrocytes through a number of receptor-ligand interactions, including the erythrocyte binding antigen 175 (PfEBA175)-glycophorin A (GPA) receptor interaction [7, 8]. Several previous studies have reported the use of antibodies against PfEBA175 to inhibit *P. falciparum* invasion of erythrocytes by blocking the PfEBA-GPA interaction [9]. High levels of PfEBA175 antibodies have also been reported to reduce *P. falciparum* parasite burden in malaria-endemic areas [10, 11].

The ookinete surface antigen, Pfs230 is found on the surface of gametocytes and thus exposed to the human immune system. Full length as well as various fragments of Pfs230 have been found to elicit humoral immune response in individuals infected with sexual stages of the parasite. Antibodies against Pfs230 have been found to prevent gamete fusion and subsequent completion of the sporogonic parasite life cycle in the mosquito [12] and serves the purpose of blocking malaria transmission.

The human host responds to the injection of mosquito saliva during a bite by producing antibodies, including antibodies against the *Anopheles gambiae* salivary protein gSG6 peptide 1 (gSG6-P1) [13, 14]. Seroprevalence and intensity of IgG responses against gSG6-P1 can be used to estimate the anopheline biting rate in malaria-endemic areas [15, 16] and can serve as a potential biomarker of human exposure to mosquito bites [17].

Rural areas have higher risks of malaria exposure than peri-urban areas due to improved socioeconomic status, housing systems, and limited mosquito breeding sites. However, the two study sites, Obom (rural area) and Simiw (peri-urban area) have similarly high malaria prevalence [11, 18]. However, the risk factors contributing to the high malaria prevalence in these communities are unknown.

The study hypothesized that there would be a high prevalence of *Plasmodium*-specific antibody responses in communities with high levels of exposure to mosquito vectors. This study compared IgG responses against PfCSP, PfEBA175 and Pfs230 in individuals with varying levels of anti-gSG6-P1 IgG levels among age groups and gender, in two malaria-endemic communities in Ghana.

## Methods

### Study site and Population

This study used samples from a study that has already been published [18]. The cross-sectional study was conducted from December 2019 to January 2020 in two malaria endemic communities in Southern Ghana. The communities include Simiw, a rural community within the Komenda Edina Eguafo Abrim District (KEEA) in the Central Region of Ghana and Obom, a peri-urban community within the Ga South Municipality of the Greater Accra Region of Ghana. Malaria transmission is similar in both Simiw and Obom is perennial with the peak malarial season occurring with the major raining season between June and August [11].

A total of 638 volunteers aged 6 months to 70 years without any symptoms of clinical malaria were recruited from Simiw (N=347) and Obom (N=291). The study was approved by the Institutional Review Board (IRB) of the Noguchi Memorial Institute for Medical Research (NMIMR no. 024/14-15 & 089/14-15). Written informed consent including assent for adolescents aged from 11 to 17, and parental consent for children aged 17 years and below were obtained from all the recruited study participants.

### Sample Collection and Processing

An earlier report has described the sample collection and processing steps involved in this study [18]. Briefly, a digital thermometer was used to measure the axillary temperature of each study participant. Subsequently, 1 ml of whole blood was collected into EDTA vacutainer tubes. Ten (10) μl aliquot of blood was used to prepare thick and thin blood smears. The remaining whole blood was separated into packed cells that were used for DNA extraction and subsequent *P. falciparum* species PCR and plasma that was stored at −20 °C for future use.

### Estimation of *Plasmodium* parasite density

The blood films were processed and stained with Giemsa according to WHO guidelines [19]. The microscopy data used in this study was obtained from the larger previously published dataset [18].

### Polymerase chain reaction detection of *Plasmodium falciparum*

The extracted genomic DNA was subjected to Nested PCR using the previously published protocol. The PCR data used in this study was obtained from the larger previously published dataset [18].

### Indirect Enzyme-Linked Immunosorbent Assay (ELISA)

The recombinant antigens Pfs230proC (pro region), PfEBA-175.5R (region III-V) and PfCSP (full length) were produced in *L. lactis* as previously described by Acquah *et al*., (2017) [20], whilst the gSG6-P1 peptide was synthesized and purified (>95%) by Genepep SA (Saint Jean de Védas, France), shipped lyophilized, resuspended in 0.22 µm ultrafiltered water and stored in aliquots at −20° C until used.

IgG antibody levels against gSG6-P1, PfCSP, Pfs230proC and PfEBA-175.5R were quantify by indirect ELISA as described by Acquah *et al*., (2017) [20]. A 96-well NUNC Maxisorp ELISA plate was coated with either 1 µg/well of gSG6-P1or CSP in phosphate buffered saline (PBS, pH 7.2), 20 µl/well EBA-175.5R in PBS, pH 7.2 or 1 µg/well of Pfs230proC in carbonate buffer, pH 9.0 and incubated overnight at 4 °C. The plates were then washed three times with 250 µl/well of wash buffer (PBS containing 0.05% Tween 20 (PBS-T)). The plates were subsequently blocked with 200 µl/well of blocking buffer (3% skimmed milk in PBS) and incubated for an hour. The plates were then washed three times and incubated for an hour with duplicate samples comprising of 100 µl/well diluted plasma (1:200), a positive control sample obtained from a pool of seropositive individuals and negative control samples obtained from various pools of seronegative individuals from Denmark. Purified polyclonal IgG (PB055, The Binding Site) at a starting concentration of 100 ng/µl was serially diluted 3-fold for 6 additional concentrations and used as a standard calibrator. The plates were washed three times and incubated for an hour with 100 µl/well of 1:3000 dilution of goat antihuman IgG-HRP. The plates were incubated with peroxidase substrate 3,3’,5,5’-teramethylbenzidine (TMB) for 10 minutes after the final washing step. 100 µl of 0.2 mM sulfuric acid was added to halt the enzymatic reactions and the optical densities (OD) of the contents in the wells were read at 450 nm using a Multiskan FC plate reader (Thermo Scientific, USA).

### Data and Statistical analyses

The optical density (OD) values obtained from the plate reader was converted to antibody concentrations using the four-parameter curve-fitting application ADAMSEL FPL (Ed Remarque^®^). A sample was defined as seropositive when the IgG concentrations obtained were higher than the average antibody concentrations of the negative control samples (malaria naïve serum samples) plus two standard deviations. All data were entered into Microsoft Excel (Microsoft Corp., Redmond, WA, USA), and statistically analyzed with GraphPad Prism software, version 9.0.2 (GraphPad Software, San Diego, CA, USA). The data were grouped into study sites, diagnostic tests, gender and age categories. The seroprevalence of gSG6-P1, CSP, Pfs230proC and EBA-175.5R IgG antibodies were determined using simple counts and proportions. The gSG6-P1, CSP, Pfs230proC and EBA-175.5R IgG antibody concentrations were Log 10 transformed. The gSG6-P1 concentration was also categorized into moderate (1-1.999 ng/mL), high (2-4.999 ng/mL) and extreme (>5 ng/mL) mosquito exposure and 0-0.999 ng/mL was considered as negative for mosquito exposure [14]. The differences in seroprevalence of the various IgG antibody concentration between the two study sites were determined using Chi-square statistics. Spearman correlation was performed to assess the association between IgG antibody levels and parasite density. The association between concentrations between the Obom and Simiw were determined using Mann Whitney U test statistics. Statistical significance was set at *P* < 0.05.

## Results

### Demographic characteristics of the study - population

A total of 639 individuals from Obom (N=292 and Simiw (N=347) were enrolled in the study (Table 1). Majority of the participants in Obom 55.1% (161/292) and Simiw 68.0% (236/347) were female. Although none of the participants presented with clinical symptoms of malaria, 39.7% (116/292) and 50.1% (174/347) of the participants from Obom and Simiw respectively tested positive for *P. falciparum* by PCR. There was similar median (range) parasite density at both Obom 440 (40-12688)/µL and Simiw 336 (32-28057)/µL (Table 1).

**Table 1.** Demographic characteristics of the study participants df, degree of freedom; P<0.05, significant; P<0.01, significant; p<0.001, highly significant

| | Study sites | | $\chi^2$ | df | p |
| --- | --- | --- | --- | --- | --- |
|  | Obom (OB) | Simiw (SW) |  |  |  |
| Sex n/N (%) |  |  |  |  |  |
| Females | 161/292 (55.1) | 236/347 (68.0) | 2.544 | 1 | 0.1107 |
| Males | 131/292 (44.9) | 111/347 (32.0) | 4.864 | 1 | 0.0274 |
| Age categories/yrs n/N (%) |  |  |  |  |  |
| <5 | 5/23 (21.7) | 17/23 (73.9) | 4.570 | 1 | 0.0328 |
| 6-10 | 69/115 (60.0) | 47/115 (40.9) | 2.566 | 1 | 0.1092 |
| 11-15 | 104/235 (44.3) | 131/235 (55.7) | 2.071 | 1 | 0.1501 |
| 16-20 | 39/69 (56.5) | 30/69 (43.5) | 0.7841 | 1 | 0.3759 |
| >20y | 75/197 (25.8) | 122/197 (61.9) | 7.523 | 1 | 0.0061 |
| Temperature/oC median (Range) |  |  |  |  |  |
| <37.5 | 36.8 (33.9-37.5) | 36.6 (34.0-37.5) | 0.02319 | 1 | 0.879 |
| >37.5 | - | 38.6 (37.6-39.3) | - | - | - |
| HB concentration g/dl |  |  |  |  |  |
| median (Range) |  |  |  |  |  |
| <10 | 9.7 (8.0-9.9) | 9.35 (7.5-9.9) | 16.09 | 1 | <0.0001 |
| 10-13 | 11.5 (10.1-12.9) | 11.3 (10.0-13.0) | 1.836 | 1 | 0.1754 |
| >13 | 14.3 (13.1-17.8) | 14.0 (31.1-17.8) | 23.16 | 1 | <0.0001 |
| Microscopic diagnosis |  |  |  |  |  |
| n/N (%) |  |  |  |  |  |
| Microscopy+ | 89/292 (30.5) | 124/347 (35.7) | 0.9891 | 1 | 0.32 |
| Microscopy- | 204/292 (69.9) | 223/347 (64.3) | 0.078 | 1 | 0.7804 |
| Parasitaemia/ul | 440.0 | 336.0 |  |  |  |
| median (Range) | (40.0-12,688.0) | (32-28,057.0) | - | - | - |
| PCR diagnosis n/N (%) |  |  |  |  |  |
| Plasmodium falciparum | 116/292 (39.7) | 174/347 (50.1) | 2.817 | 1 | 0.0933 |
| Plasmodium malariae | 42/292 (14.4) | 62/347 (17.9) | 1.02 | 1 | 0.3126 |
| Plasmodium ovale | 17/292 (5.8) | 25/347 (7.2) | 0.4333 | 1 | 0.5104 |

### gSG6-P1

Exposure to *Anopheles* bites was evaluated by quantifying IgG levels specific to the gSG6-P1 salivary peptide from *An. gambiae*. Overall, the participants from Simiw had significantly (Mann Whitney U=38201, p<0.0001) higher median gSG6-P1 IgG concentration compared to participants from Obom (S1 Figure). Participants from Simiw within the 11-15 years old age group had significantly (p<0.0001) higher levels of gSG6-P1 IgG relative to similarly aged children from Obom. All other age matched categories in Obom and Simiw had similar median gSG6-P1 IgG levels (Figure 1). Participants from Simiw with PCR confirmed *P. falciparum* infection had higher gSG6-P1 IgG antibodies (median (95% CI) ng/mL) (2.590 (2.452-2.783) ng/mL) than their counterparts in Obom (2.119 (1.957-2.345) ng/mL), p<0.0001 (Figure 2 **a**). Although there was no significant difference in anti-gSG6-P1 IgG levels between individuals with or without PCR confirmed *P. falciparum* infections at Obom (p=0.2800), there was a significant difference between individuals with and without PCR detectable *P. falciparum* in Simiw (p=0.0170) (Figure 2a).

**Figure 1.**
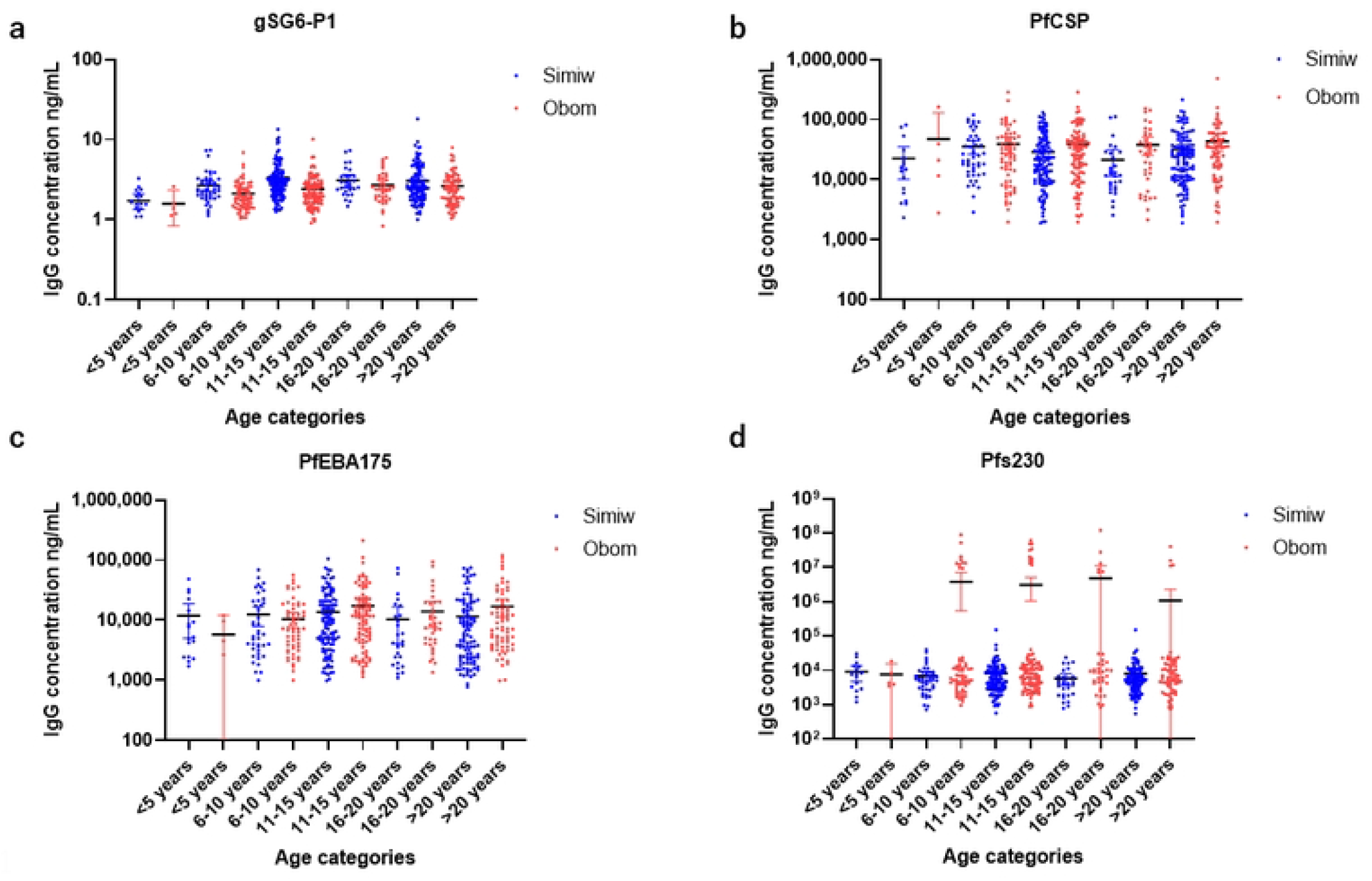
Age stratified IgG levels. Each dot presents the individual immunoglobulin G (IgG) antibodies concentration among age categories of the study participants between Obom (OB) and Simiw (SW) against **a** Anopheles salivary gSG6-P1; **b** P. *falciparum* CSP; **c** *P. falciparum* erythrocyte binding antigen-175 (PfEBA-175); **d** gametocyte surface antigen Pfs230. The horizontal bar indicates the median value of specific IgG levels.

**Figure 2.**
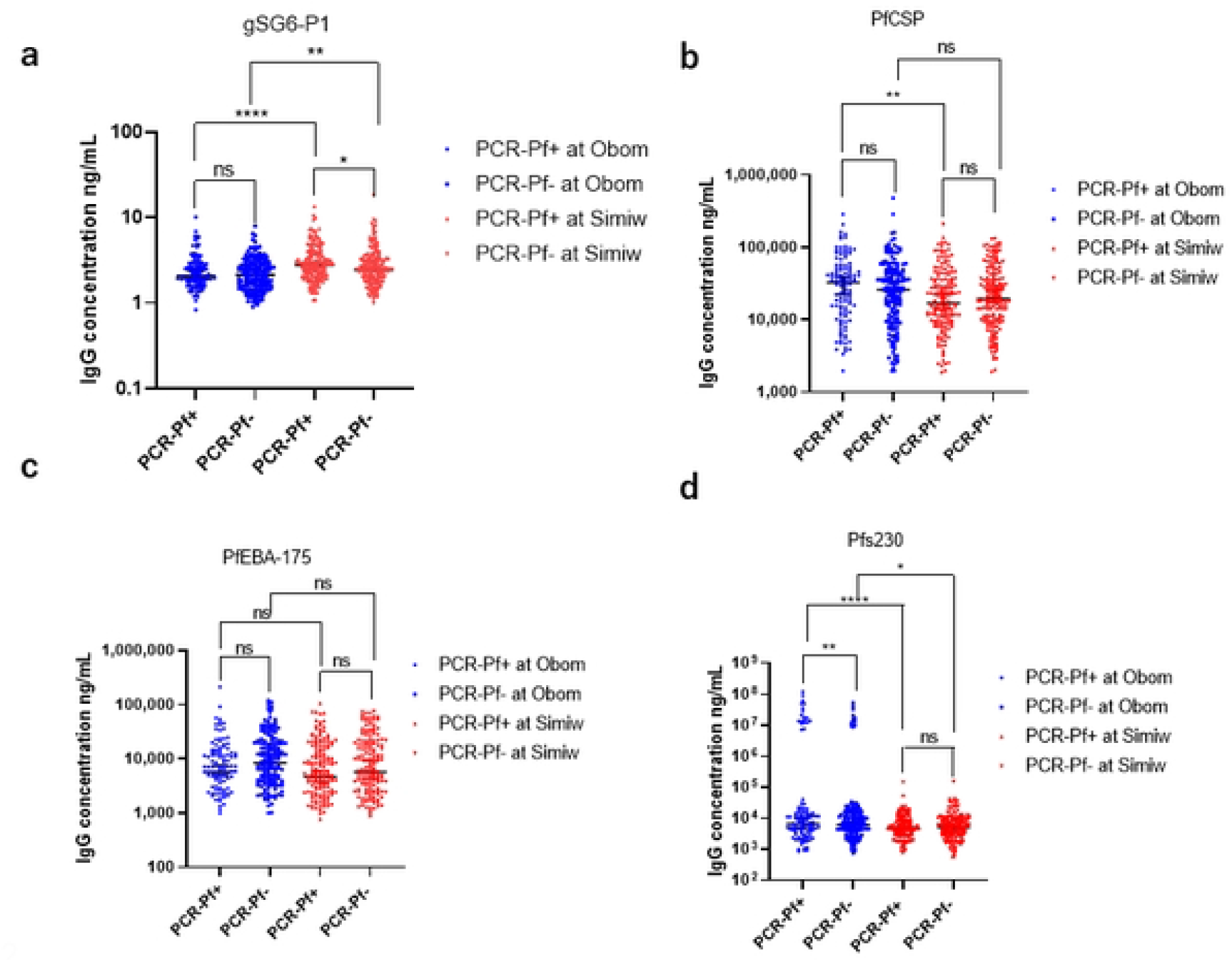
Comparison of IgG levels in PCR-confirmed *P. falciparum* positive and negative individuals. **a** The distribution of Anopheles salivary gland gSG6-P1; **b** *P. falciparum* CSP; **c** *P. falciparum* erythrocyte binding antigen-175 (PfEBA-175); **d** gametocyte surface antigen Pfs230 immunoglobulin G (IgG) antibodies concentration in the PCR confirmed *P. falciparum* infections and uninfected case between Obom (OB) and Simiw (SW) study communities The significance was tested using Mann Whitney U test; ns (p>0.05) not significant; * (P<0.05), significant; ** (P<0.01), significant; ***(p<0.001), highly significant; **** (p<0.0001), highly significant

Highly significant difference (χ2=19.55, df=1, p<0.0001) was observed in the seroprevalence of gSG6-P1 IgG in Simiw (81.3% (282/347)) than Obom (65.5% (190/292)) (Table 2). A similar observation was made for uninfected individuals (p=0.0027) as well as after comparing male (p=0.0001) and female (p=0.002) participants from Obom and Simiw (Figure S2 **a**). However, the seroprevalence of gSG6-P1 IgG amongst females in Obom (71.4% (115/161)) was similar to females in Simiw (83.5% (197/236)), and males in Simiw (76.6% (85/111)) than in Obom (57.3% (75/131)). Also, there was no significant difference across the age categories between Obom and Simiw (Table 2).

**Table 2:** Seroprevalence of IgG antibodies in Obom and Simiw df, degree of freedom; P<0.05, significant; P<0.01, significant; p<0.001, highly significant

| Variables | gSG6-P1 IgG |  |  |  |  | PfCSP IgG |  |  |  |  | PfEBA-175.5R IgG |  |  |  |  | Pfs230proC IgG |  |  |  |  |
| --- | --- | --- | --- | --- | --- | --- | --- | --- | --- | --- | --- | --- | --- | --- | --- | --- | --- | --- | --- | --- |
| | Obom | Simiw | $\chi^2$ | df | p | Obom | Simiw | $\chi^2$ | df | p | Obom | Simiw | $\chi^2$ | df | p | Obom | Simiw | $\chi^2$ | df | p |
| Community, n/N (%) | 190/292<br>(65.5) | 282/347<br>(81.3) | 19.55 | 1 | <0.0001 | 219/292<br>(75%) | 234/327<br>(71.6) | 0.1439 | 1 | 0.7044 | 266/292<br>(91.1%) | 304/327<br>(93.0) | 0.0305 | 1 | 0.8613 | 244/292<br>(83.6) | 275/327<br>(84.1) | 0.0029 | 1 | 0.9572 |
| Sex, n/N (%) |  |  |  |  |  |  |  |  |  |  |  |  |  |  |  |  |  |  |  |  |
| Male | 75/131<br>(57.3) | 85/111<br>(76.6) | 0.3635 | 1 | 0.5466 | 95/131<br>(72.5) | 83/111<br>(74.8) | 0.0239 | 1 | 0.8771 | 121/131<br>(92.4) | 97/111<br>(87.4) | 0.0872 | 1 | 0.7678 | 110/131<br>(84.0) | 82/111<br>(73.9) | 0.4328 | 1 | 0.5106 |
| Female | 115/161<br>(71.4) | 197/236<br>(83.5%) | 1.003 | 1 | 0.3165 | 124/161<br>(77.0) | 160/236<br>(67.8) | 0.6571 | 1 | 0.4176 | 145/161<br>(90.1) | 207/236<br>(87.7) | 0.0315 | 1 | 0.8591 | 134/161<br>(83.2) | 193/236<br>(81.8) | 0.0134 | 1 | 0.9079 |
| $\chi^2$ | 1.366 | 0.2473 | | | | 0.1117 | 0.3043 | | | | 0.022 | 0.0005 | | | | 1.666 | 32.0 | | | |
| df | 1 | 1 |  |  |  | 1 | 1 |  |  |  | 1 | 1 |  |  |  | 1 | 1 |  |  |  |
| p | 0.2425 | 0.619 |  |  |  | 0.0945 | 0.5812 |  |  |  | 0.882 | 0.982 |  |  |  | 0.1967 | <0.0001 |  |  |  |
| Age<br>categories, n/N<br>(%) |  |  |  |  |  |  |  |  |  |  |  |  |  |  |  |  |  |  |  |  |
| <5 yr | 1/5<br>(20.0) | 9/17<br>(52.9) | 0.731 | 1 | 0.3926 | 4/5<br>(80.0) | 11/17<br>(64.7) | 0.0752 | 1 | 0.7839 | 5/5<br>(100.0) | 17/17<br>(100.0) | 0 | 1 | >0.9999 | 5/5<br>(100.0) | 14/17<br>(82.4) | 0.0712 | 1 | 0.7896 |
| 6-10 yr | 36/68<br>(52.9) | 39/47<br>(83.0) | 2.27 | 1 | 0.1319 | 48/68<br>(70.6) | 40/47<br>(85.1) | 0.4279 | 1 | 0.513 | 61/68<br>(89.7) | 42/47<br>(89.4) | 0.0002 | 1 | 0.9889 | 54/68<br>(79.4) | 36/47<br>(76.6) | 0.0158 | 1 | 0.8998 |
| 11-15 yr | 69/104<br>(66.3) | 113/130<br>(86.9) | 1.798 | 1 | 0.1799 | 79/104<br>(76.0) | 87/130<br>(66.9) | 0.3872 | 1 | 0.5338 | 94/104<br>(90.4) | 113/130<br>(86.9) | 0.0415 | 1 | 0.8387 | 94/104<br>(90.4) | 102/130<br>(78.5) | 0.5304 | 1 | 0.4665 |
| 16-20 yr | 31/39<br>(79.5) | 28/30<br>(93.3) | 0.2032 | 1 | 0.6521 | 28/39<br>(71.8) | 17/30<br>(56.7) | 0.3653 | 1 | 0.5456 | 33/39<br>(84.6) | 27/30<br>(90.0) | 0.0301 | 1 | 0.8622 | 31/39<br>(79.5) | 21/30<br>(70.0) | 0.1164 | 1 | 0.733 |
| > 20 yr | 53/76<br>(69.7) | 93/123<br>(75.6) | 0.1284 | 1 | 0.7201 | 60/76<br>(78.9) | 88/123<br>(71.5) | 0.1966 | 1 | 0.6576 | 73/76<br>(96.1) | 105/123<br>(85.4) | 0.3127 | 1 | 0.576 | 60/76<br>(78.9) | 102/123<br>(82.9) | 0.0506 | 1 | 0.822 |

### PfCSP

Participants from Obom had significantly higher IgG levels against the CSP (27781 (23674-33921) ng/mL) compared to participants from Simiw (18330 (15984-22924) ng/mL) (p=0.002, S1 Figure **b**). With exception of the 16–20-year age category (p=0.049), all other age categories in both Obom and Simiw had similar anti-CSP IgG levels (Figure 1 **b**). No significant difference was observed in the median anti-CSP IgG in *P. falciparum* PCR positive and negative participants at Obom (p=0.1868) and Simiw (p=0.3736). However, *P. falciparum* PCR-positive individuals in Obom had significantly (p=0.0015) higher median anti-CSP IgG relative to counterparts in Simiw (Figure 2 **b**). The males and females at both Obom (p=0.1804) and Simiw (p=0.7798) had similar anti-CSP IgG levels (S2 figure **b**).

Overall, seroprevalence of anti-CSP IgG was similar in Obom and Simiw (p=0.7044) and not influenced by gender (Table 2).

### PfEBA-175.5R

There was no significant difference in IgG responses against the erythrocytic binding antigen, PfEBA-175.5R in total population and across the various age categories in individuals from both Simiw and Obom (p > 0.05) (Figure 1 **c**, S1 Figure **c**). This observation is confirmed by similar parasite prevalence detected by both microscopy and PCR in Obom 30.6% (89/291) and 39.5% (115/291) respectively and (35.7% (124/347) vs 50.1% (174/347) respectively in Simiw (Table 1). In addition, parasite densities in samples from Obom, 440.0 (40.0-12,688.0) and Simiw, 366.0 (32.0-28,052.0) were similar (Table 1). There was no significant difference in the PfEBA-175.5R seroprevalence in Obom (91.1% (266/292)) and Simiw (93% (304/327)) (χ2 =0.0305, df=1, p=0.8613) overall. Gender did not have any influence on the level and the seroprevalence of anti-PfEBA-175.5R IgG in Obom (p=0.882) and Simiw (p=0.982) (Table 2; S2 figure **c**).

### Pfs230proC

Exposure to gametocytes was determined by measuring antibody responses against the gametocyte and ookinete surface antigen Pfs230proC. Significantly (p=0.0006) higher IgG responses against Pfs230 were measured in participants from Obom (5993 (5112-7296) ng/mL) compared to Simiw (4822 (4306-5328) ng/mL) (S1 Figure **d**). There was no significant difference in the median concentration of Pfs230proC IgG among the various age categories in Obom and Simiw, except among the 11-15 years (p=0.007) and 16-20 years (p=0.0477) old age groups, where participants from Obom had significantly higher levels than their counterparts from Simiw (Figure 1 **d**). Although there was no significant difference in Pfs230proC IgG levels between individuals that tested negative or positive for *P. falciparum* by PCR in Simiw (p =0.240), there was a significant difference in Obom (p =0.0071). Individuals with PCR confirmed infections in Obom had significantly higher Pfs230proC IgG levels than individuals with PCR confirmed infections in Simiw (p =0.003) (Figure 2 **d**). Similarly, the uninfected individuals in Obom had significantly higher Pfs230proC IgG levels than uninfected individuals at Simiw (p=0.048) (Figure 2 **d**). There was however no significant difference in the median concentration of Pfs230proC IgG antibodies between males and females at Obom p=0.353 and at Simiw p=0.424 (S2 Figure **d**). The median concentration of Pfs230proC IgG in males from Obom (6562 (5021-8994) was significantly (p=0.003) higher than detected in males from Simiw (4269 (3481-5328) (S2 Figure **d**). However, there was no significant (p=0.053) difference in Pfs230proC IgG in females from Obom and Simiw. The seroprevalence of anti-Pfs230proC IgG in Obom 83.6% (244/292) and Simiw 84.1% (275/327) was similar χ2=0.0029, df=1, p=0.9572 (Table 2).

### Association between IgG concentration and parasite density

Spearman correlation showed no significant correlation between IgG antibody concentrations of gSG6-P1, PfCSP, Pfs230proC and PfEBA-175.5R and parasite density (Figure 3 **a-d**). The correlation coefficient (95% CI) of association between gSG6-P1 IgG antibody and parasite density was r= 0.03565 (−0.2785 to 0.2492), p =0.7416 at Obom and r =−0.0968 (−0.2785 to 0.09163), p =0.2992 at Simiw (Figure 3 **a**).

**Figure 3.**
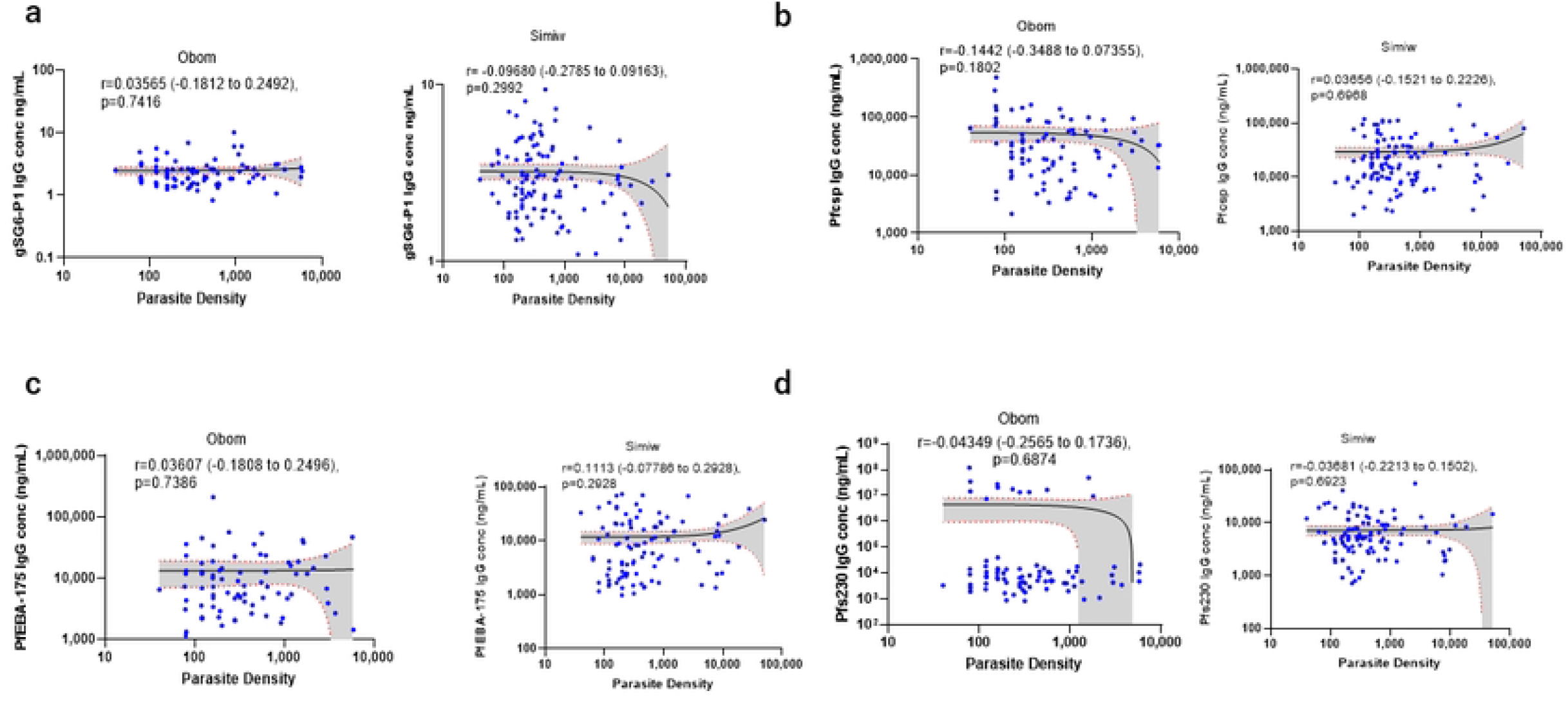
Spearman correlation between the distribution of IgG levels and *P. falciparum* parasite density. **a** Spearman correlation between the distribution of Anopheles salivary gland gSG6-P1; **b** *P. falciparum* CSP; **c** *P. falciparum* erythrocyte binding antigen-175 (PfEBA-175); **d** gametocyte surface antigen Pfs230 immunoglobulin G (IgG) antibodies and the distribution of *P. falciparum* parasite density in the study communities. **The X- and Y-axis are represented in log10. The significance (p>0.05) not significant; (P<0.05), significant; (P<0.01), significant; (p<0.001), highly significant; (p<0.0001), highly significant

### Association between salivary gland IgG levels and *P. falciparum* IgG levels

The PfCSP, PfEBA-175 and Pfs230 IgG antibody concentrations were compared across Obom and Simiw among individuals with similar mosquito exposure levels (gSG6-P1 IgG concentrations of 1-1.999 ng/mL, 2-4.999 ng/mL and >5 ng/mL) (Fig 4). The results show significantly lower levels of anti-PfCSP IgG (sporozoite exposure) among subjects with moderate 1-1.999 ng/mL IgG gSG6-P1 antibody level (Mann-Whitney U=4128, p=0.002) at Simiw relative to Obom. However, there was no significant difference in anti-Pfs230 and PfEBA-175 IgG concentration among individuals in both Obom and Simiw with moderate gSG6-P1 IgG responses (1-1.999 ng/mL) (Figure 4 **a**). Individuals with high gSG6-P1 IgG concentrations (2-4.999 ng/mL) in Simiw had significantly lower IgG concentration of Pfs230 (Mann-Whitney U=11740, p=0.0004), and PfEBA-175 (Mann-Whitney U=27103, p=0.0468) compared to individuals from Obom with similar gSG6-P1 IgG responses. However, IgG responses against PfCSP in individuals from Obom and Simiw were similar in people high levels (2-4.999 ng/mL) of anti-gSG6-P1 IgG (Figure 4 **b**).

**Figure 4.**
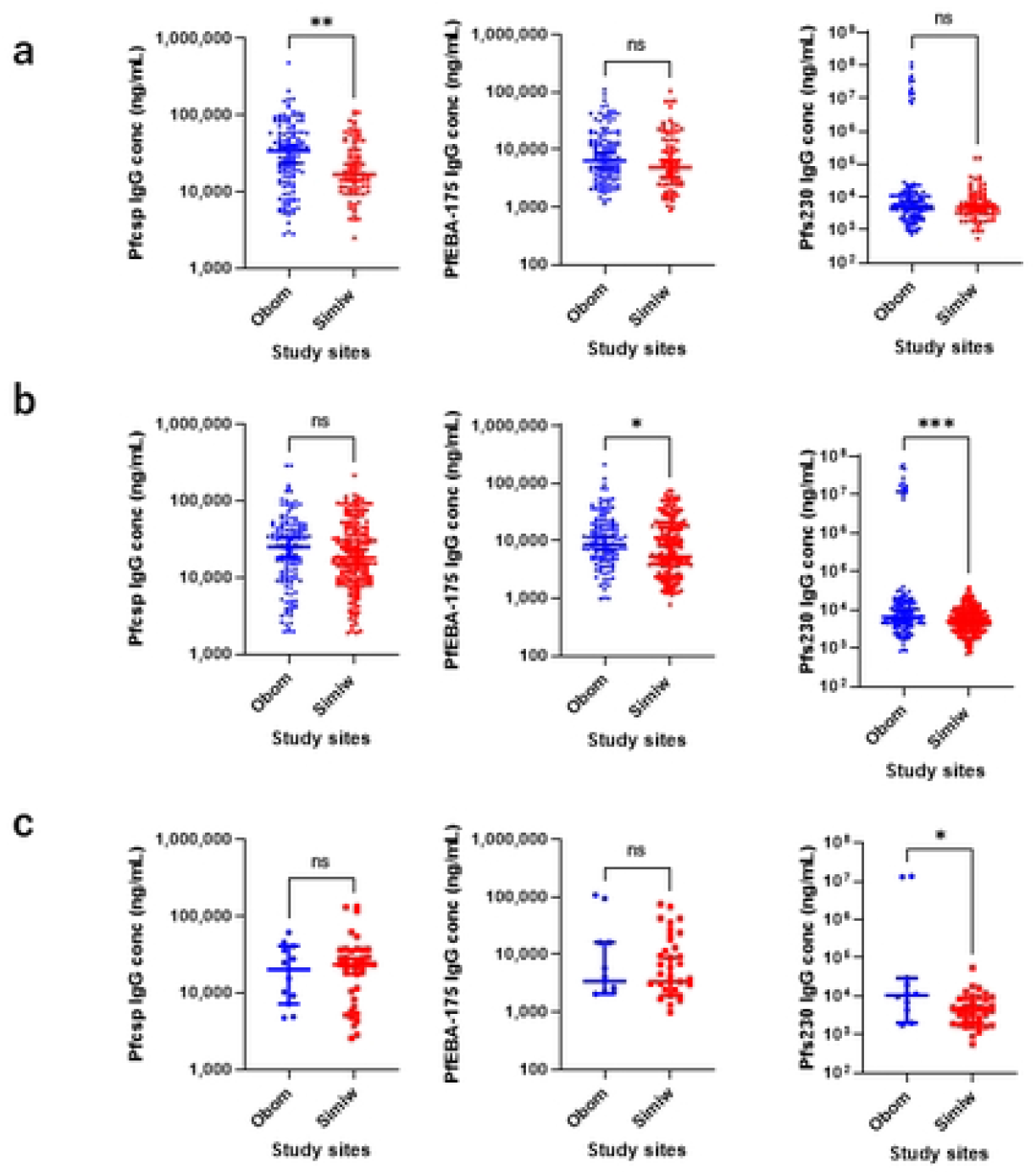
*P. falciparum* stage-specific IgG levels in individuals with similar mosquito exposure. Association of IgG antibodies of PfCSP, PfEBA-175, and Pfs230 between Obom and Simiw with **a** 1-2 ng/mL; **b** 2-5 ng/mL; **c** > 5 ng/mL gSG6-P1 IgG antibody exposure

Individuals with >5 ng/mL anti-gSG6-P1 IgG levels in Obom had significantly higher levels Pfs230 (Mann-Whitney U=146.5, p=0.034) compared to individuals from Simiw with similar anti-gSG6-P1 IgG levels (Figure 4 **c**). The significant difference in Pfs230 IgG antibodies between Obom and Simiw was due to a small sub-group of individuals in Obom who had high IgG antibodies of Pfs230 (S3 Figure **a** & **b**).

The correlation between levels of IgG antibodies to PfCSP, Pfs230 and PfEBA-175 to gSG6-P1 IgG were assessed among individuals within the three gSG6-P1 IgG categories; 1-1.999 ng/mL, 2-4.999 ng/mL and >5 ng/mL gSG6-P1 IgG concentrations (S4 fig). There was a significant negative correlation between PfEBA-175 IgG and individuals with moderate gSG6-P1 IgG (1-1.999 ng/mL) in Simiw (r=−0.2702 (−0.4643 to −0.05156), p=0.014) but not in Obom (r=−0.0738 (−0.2452 to 0.102), p=0.397). No association was observed between anti-Pfs230 or PfCSP IgG levels and moderate (1-1.999 ng/mL) IgG gSG6-P1 levels.

A similar significantly negative correlation was identified between high (2-4.999 ng/mL) gSG6-P1 IgG and anti-Pfs230 IgG levels in Simiw (r=−0.1357 (−0.267 to 0.003346), p=0.049) but not in Obom (r=−0.1132 (−0.2768 to 0.05682), p=0.178). No other associations were identified between moderate and high gSG6-P1 IgG and the measured antiparasite IgG in Simiw and Obom (S4 Figure **a & c**).

## Discussion

Several factors regulate the dynamics of *P. falciparum* infection and transmission in malaria-endemic settings. The gold standard for measuring malaria transmission intensity is using entomological inoculation rate (EIR), however, the insufficiencies associated with the traditional estimation of EIR using infected mosquitoes have led to the identification of some serological markers of exposure to malaria parasites as alternative and more effective measures of malaria transmission intensity in both high and low transmission settings [14, 15].

Variations in host factors and exposures to malaria parasites can impact the immune responses generated by the host [21]. Host-specific immune response to malaria infection can offer protection against severe malaria and control transmission in endemic areas [20]. There is still however, gaps in our understanding of the role of antibodies against various malaria parasite stage or mosquito antigens play in influence transmission.

This study assessed antibody levels against the sporozoite stage of the malaria parasite, PfCSP, the asexual blood stage parasites, PfEBA175 and the sexual blood stage parasite Pfs230 in individuals with varying levels of exposure to the mosquito vector, in two communities of high malaria transmission in southern Ghana.

The formulation of malaria control and elimination programmes are based on the intensity of malaria transmission within a community. The IgG to gSG6-P1 is a proxy to exposure and in some contexts has been associated to clinical outcomes [15, 17]. It could be used in assessing temporal and spatial variations in exposure to Anopheles mosquito bites [14, 17]. The study showed a significantly high seroprevalence and high median concentration of gSG6-P1 IgG antibodies in Simiw compared to Obom, which could pose a high risk of increased malaria transmission in Simiw. The higher exposure to mosquitoes in Simiw is likely due to the influence of environmental factors such as having large areas of stagnant water, which serves as mosquito breeding sites. Also, individuals from Obom are more urban and likely to protect themselves from mosquito bites [22, 23]. The increased levels of IgG to gSG6-P1 in females at both study sites was surprising as it has previously been suggested that certain behavior of males, including staying outdoors after dark predisposes them to more frequent encounters with mosquitoes [24]. The high exposure of females to mosquitoes could be a result of changes in the behavior of mosquitoes, where there are increased reports of mosquitoes biting during the early morning or an increase in biting indoor [25-27], when females are busy sweeping and doing other household chores.

= Significantly higher levels of PfCSP IgG in Obom compared to Simiw indicates higher vector competency and effective malaria transmission [28, 29] at Obom than in Simiw. The likely explanation could be that mosquitoes in Obom are more efficient at transmitting malaria, which cannot be validated as we did not obtain information on the mosquito population in Obom and Simiw. Another explanation could be that there were more transmission reservoirs (gametocyte carriers) in Obom relative to Simiw where there was an abundance of mosquitoes. Indeed, participants from Obom presented higher anti-Pfs230 levels.

High IgG antibodies to PfEBA 175 in people living in *P. falciparum* high endemic communities, protect against erythrocytic stage invasion and symptomatic malaria [10, 11]. There was no significant difference in the median concentration and seroprevalence of IgG antibodies to PfEBA 175 in individuals from Simiw and Obom. The observation was expected as the two communities had similar malaria prevalence and parasite densities.

Gametocyte carriage is essential for malaria transmission however, exposure to gametocytes can reduce malaria transmission as some antibodies against gametocyte antigens, including antibodies against specific regions of Pfs230 serve as transmission blocking antibodies, which can prevent the development of the parasite in the mosquito [12, 30]. The study observed that although the seroprevalence of IgG antibodies against Pfs230 was very similar in Obom and Simiw, there was a significantly higher median concentration of IgG antibodies to Pfs230 in Obom compared to Simiw. Since specific Pfs230 IgG antibodies can be transmission blocking [12, 36] and or serve as a marker of exposure to gametocytes. It was observed that a sub-population of participants from Obom had very high levels of IgG antibodies, suggesting a recent exposure to high gametocyte densities. Also, a significant number of this sub-population were confirmed to harbor *P. falciparum*.

The study compared similar levels of exposure to mosquitoes in Obom and Simiw; the result showed that at a moderate concentration of gSGP1 IgG (concentration of 1-1.999 ng/mL), Simiw had lower exposure to *P. falciparum* sporozoites, indicating a higher prevalence of uninfected mosquitoes than Obom. Exposure to uninfected mosquito bites has been suggested to reduce liver stage hypnozoites as well as asexual stage parasite burden relative to exposure to malaria-infected mosquitoes [37, 38]. A previous report also identified an increase in interleukin-12, gamma interferon and inducible nitric oxide synthase after repeated exposure to uninfected mosquito bites. These T-helper 1 (Th1) immunity regulate asexual stage parasitaemia in murine model [39]. This study showed no significant difference in PfEBA-175 IgG antibodies in Obom and Simiw, suggesting similar exposure to asexual parasites. Interesting, individuals in Obom and Simiw with high (2-5 ng/mL) levels of gSGP1 IgG antibodies (high exposure to mosquito bites) had similar exposure to sporozoites (PfCSP IgG antibodies). However, Simiw had lower asexual stage parasite density and exposure to asexual parasites (PfEBA-175 IgG antibody levels) compared to Obom, supporting the observation that mosquito salivary components reduce asexual parasite burden [37,38].

In conclusion, malaria transmission dynamics is highly complex. The similar malaria transmission intensity identified in the two communities resulted from a different combination of vector, environmental and host factors. With one community likely having a higher prevalence of uninfected mosquitoes and the other a larger reservoir of gametocyte carriers.

## Acknowledgement

We thank all the study participants for willingly participating in the study

## Funding

This project was partially funded by the JEAI-STIMULI project awarded to LEA; AKK was partly supported with the 2021 Ghana government books and research allowance;

## Author’s contribution

Conceptualization; LEA

Formal analysis; KKA, LEA, SSK, AP

Methodology; SSK

Supervision; LEA, FP

Validation; KKA, SSK, RT, KAK

Visualization; SSK, KKA

Writing – original draft; KKA, SSK, LEA,

Writing – review & editing; KKA, LEA, AP, RT, NN, RT, KAK

## Competing interests

The authors have nothing to declare

## Table and Figure legends

S1 Figure: The overall **Immunoglobulin G (IgG) antibody levels among the study communities**

**a** Distribution of Anopheles salivary gland gSG6-P1; **b** *Plasmodium falciparum* CSP; **c** *Plasmodium falciparum* EBA 175; **d** *Plasmodium falciparum* Pfs230 immunoglobulin G (IgG) antibodies among the study communities

The significance was tested using Mann Whitney U test; ns (p>0.05) not significant; * (P<0.05), significant; ** (P<0.01), significant; ***(p<0.001), highly significant; **** (p<0.0001), highly significant

S2 Figure: **Immunoglobulin G (IgG) levels in male and female study participants**

**a** The Anopheles salivary gland; **b** The *P. falciparum* CSP; **c** The *P. falciparum* erythrocyte binding antigen-175 (PfEBA-175); **d** The gametocyte surface antigen Pfs230 immunoglobulin G (IgG) antibodies concentration among male and female study participants between Obom (OB) and Simiw (SW)

S3 Figure ***P. falciparum* stage-specific IgG antibodies with similar mosquito exposures after excluding sub-groups expressing high Pfs230 IgG antibody** Association of IgG antibodies of PfCSP, Pfs230, PfEBA-175 between Obom and Simiw with **a** 2-5 ng/mL; **b** > 5 ng/mL gSG6-P1 IgG antibody exposure after excluding sub-groups expressing high Pfs230 IgG antibody at Obom

S4 Figure **Correlation of *P. falciparum* stage-specific IgG antibodies across individuals with different levels of mosquito exposure**

Correlation between **a** 1-2 ng/mL; **b** 2-5 ng/mL; **c** > 5 ng/mL gSG6-P1 IgG antibodies and PfCSP, Pfs230, and PfEBA-175 in Obom and Simiw

## Notes

### Competing Interest Statement

The authors have declared no competing interest.

